# Investigating on relationship between effective quantum efficiency and irradiance

**DOI:** 10.1101/222414

**Authors:** Zi-Piao Ye, Shuang-Xi Zhou, Xiao-Long Yang, Hua-Jing Kang, Piotr Robakowski

## Abstract

**Abstract:** Models describing the relationship between effective quantum efficiency of PS II (*Φ*_PSII_) and irradiance (*I*) are routinely used to determine how irradiance influences effective quantum efficiency and photosynthetic electron transport rate (*ETR*). However, with no single model one can accurately describe the relationship between *Φ*_PSII_ and I, and explain the interdependence between *Φ*_PSII_ and biophysical properties of photosynthetic pigments, especially in plants growing under low level irradiances. Basing on the mechanistic model of photosynthetic electron transport rate we have developed the model of the relationship between *Φ*_PSII_ and *I*. The new model reveals that *Φ*_PSII_ increases with photochemistry (*k*_P_) and heat dissipation (*k*_D_). Furthermore, the values of key parameters calculated using the new model were compared with the values calculated with two other empirical models. The new model was perfectly fitted to the light-response curves of *Φ*_PSII_. The key calculated photosynthetic parameters: maximum *Φ*_PSII_, maximum *ETR* and their corresponding saturation irradiance were close to the measured values. In addition, our model associates *Φ*_PSII_ with intrinsic features of photosynthetic pigments. We concluded that *Φ*_PSII_ decreased with increasing *I* due to the decrease in the effective absorption cross-section of photosynthetic pigments molecules.

**Highlight:** A model of the relationship between effective quantum efficiency of PS II (*Φ*_PSII_) and irradiance (*I*) has been developed. Using this new model it was found that *Φ*_PSII_ decreased with increasing *I* due to the decrease in the effective absorption cross-section of photosynthetic pigments molecules.

**Abbreviations:**

*ETR*: Electron transport rate
*ETR*_max_: Maximum electron transport rate
*F*: Steady-state fluorescence
*F*_m′_: Maximum fluorescence in the light
*F*_v_: Variable fluorescence yield of the dark-adapted leaf
*g*_i_: Degeneration of energy level of photosynthetic pigment molecules in the ground state *i*
*g*_k_: Degeneration of energy level of photosynthetic pigment molecules in the excited state *k*
*I*: Irradiance
*NPQ*: Non-photochemical quenching
*N*_0_: Total light-harvesting pigment molecules
*PAR*_sat_: Saturation irradiance corresponding to *ETR*_max_
*k*_P_: Rate of photochemical reaction
*k*_D_: Rate of non-radiative heat dissipation
PS II: Photosystem II
*a*_e_: Initial slope of light-response curve of electron transport rate
*α′*: Fraction of light absorbed by PS II
*β′*: Leaf absorptance
*ξ*_1_: Probability of photochemistry
*ξ*_2_: Probability of non-radiative heat dissipation
*ξ*_3_: Probability of fluorescence
*σ*_ik_: Eigen-absorption cross-section of photosynthetic pigment from ground state *i* to excited state *k* due to light illumination
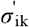: Effective optical absorption cross-section of photosynthetic pigment molecule from ground state *i* to excited state *k* due to light illumination
*φ*: Exciton-use efficiency in PS II
*τ*: Average lifetime of the photosynthetic pigment molecules in the lowest excited state
*Σ*_PSII_: Effective quantum efficiency of PS II

## Introduction

Light reactions of photosynthesis have been characterized by means of using the measurements of chlorophyll fluorescence as a useful and informative indicator (Krause & Weis, 1984; van Kooten & Snel, 1990; Rascher *et al*., 2000; van der Tol *et al*., 2014; Kalaji *et al*., 2016). The key photosynthetic parameters are: maximum quantum efficiency of photosystem II (*F*_v_/*F*_m_), effective quantum efficiency of photosystem II (*Φ*_PSII_), photosynthetic electron transport rate (*ETR*) and other chlorophyll fluorescence parameters (e.g. non-photochemical quenching, *NPQ*, coefficient of photochemical quenching, *qL*). Genty *et al*. (1989) proposed that *Φ*_PSII_ at the steady state could be calculated from the ratio of the variable to maximum fluorescence in the light. It is a breakthrough concept of using fluorescence and irradiance (*I*) or the flux of photosynthetically active radiation (*PAR*) absorbed by leaf to estimate *ETR*, which is the most widely used to assess the efficiency of plants photochemistry in different environments (Genty *et al*., 1989; Majláth *et al*., 2016; Moin *et al*., 2016). *Φ*_PSII_ represents the proportion of photons of incident light that are actually used to drive photochemistry (Maxwell & Johnson, 2000). *Φ*_PSII_ is directly associated with *ETR* (=*α*′ × *β*′ × *Φ*_PSII_ × *I*, where *α*′ is a distribution coefficient of absorption of the light energy by PS II and PS I, *β*′ is leaf absorptance, *I* is irradiance) ( Krall & Edward, 1992). Under controlled conditions, this parameter linearly relates with quantum efficiency of CO_2_ assimilation (Genty *et al*., 1989). However, in natural environment where stressors are likely to affect photosynthesis alternative processes to CO_2_ assimilation, such as photorespiration or Mehler reaction may cause discrepancy between *Φ*_PSII_ and CO_2_ assimilation (Fryer *et al*., 1998).

According to the definition of 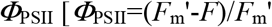, where 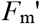 is maximum fluorescence in the light, *F* is steady-state fluorescence], it is clear that *Φ*_PSII_ is closely linked with the closure and opening of PS II in photosynthetic primary reactions as well as fluorescence emission of light-harvesting pigment molecules because chlorophyll fluorescence mainly come from these photosynthetic pigments (Baker, 2008). Photosynthesis research has long focused on *Φ*_PSII_ and *ETR* due to photosystem II (PS II) given its core role in photosynthesis, but also since PS II activity can be conveniently assayed via bio-optical techniques, e.g. chlorophyll fluorescence (e.g. Buckley & Farquhar, 2004; Robakowski, 2005; Baker, 2008; Suggett *et al*., 2010; Pavlovič *et al*., 2011). Numerous studies have used these fluorescence techniques to determine *Φ*_PSII_ and *ETR* (see the references above) and found *Φ*_PSII_ to decrease nonlinearly with increasing irradiance (Robakowski, 2005; Pavlovič *et al*., 2011; van der Tol *et al*., 2014; Córdoba *et al*., 2016). However, no single model is currently able to describe satisfactorily the relationship between *Φ*_PSII_ and *PAR*, and reveal the action of intrinsic characteristics of light-harvesting pigments in *Φ*_PSII_.

In photosynthesis, antenna pigment molecules absorb light energy to change the state of the pigments from the ground state to the excited state. Then, the excitation energy is mainly used by three inter-competing paths, i.e. photochemistry, heat dissipation and chlorophyll fluorescence emission (Müller *et al*., 2001; Oxborough, 2004; Baker, 2008). This competition among three de-excited paths (Oxborough, 2004; Baker, 2008) directly affects photosynthetic electron transport rates and formation of assimilation force (i.e. NADPH and ATP). Furthermore, light energy absorption, quantum state change, exciton resonant transfer among antenna pigment molecules and fluorescence emission in primary reaction are determined by intrinsic characteristics of the antenna pigment molecules (Govindjee, 2002; Baker, 2008; Richter *et al*., 2008 Panitchayangkoon *et al*., 2010; Sarovar *et al*., 2010). However, quantifying the nature of processes from light absorption to water splitting is still extremely limited but remains a key goal to improve models for predicting values of *Φ*_PSII_ from only light absorption measurements (Renk *et al*., 2000; Buckley & Farquhar, 2004).

As the research shows, methodologically, *Φ*_PSII_ can be directly modelled by several empirical models (Laws *et al*., 2002; Smyth *et al*., 2004; Ritchie, 2008; Ritchie & Bunthawin, 2010; Silsbe & Kromkamp, 2012) and is considered as an equivalent to *ETR*. Although these models can fit well light-response curves of electron transport rate (*ETR-I*) and light-response curves of effective quantum efficiency of PS II (*Φ*_PSII_-*I*) under normal conditions, models simulating *ETR-I* and *Φ*_PSII_-*I* in plants under low light intensities in greenhouse or under a forest trees’ canopy are yet to be developed. Modelling of these curves under such irradiance environmental conditions is necessary for the evaluation of the generality of the models. Moreover, no previous model neither explains the reasons why *Φ*_PSII_ decreased nonlinearly with increasing irradiance (Robakowski, 2005; Pavlovič *et al*., 2011; van der Tol *et al*., 2014), nor helps to understand the way intrinsic characteristics of light-harvesting pigment molecules affect *Φ*_PSII_ and determine light harvesting, light energy conversion as well as subsequent productivity (*ETR*) through PS II.

In the present study, we hypothesized that: (1) *Φ*_PSII_ should be closely related to the intrinsic properties of light-harvesting pigment molecules, closure and opening of PS II and photosynthetic enzyme kinetics of plants, (2) *Φ*_PSII_ depends on the competition of three paths of de-excitation, thus if more excitation energy is distributed to photochemistry, and less to heat dissipation, or chlorophyll fluorescence emission, *Φ*_PSII_ will be higher, and (3) behavior of the effective light absorption cross-section of light-harvesting pigment molecules will determine changes of *Φ*_PSII_.

In this study, we firstly developed a model of the relationship between *Φ*_PSII_ and *I* based on fundamental properties of light absorption and transfer of energy to the reaction centers via photosynthetic pigment molecules (see Ye *et al*. 2013a, 2013b). Firstly, here we adopt a novel approach to determine the interdependence between the properties of photosynthetic pigments and *Φ*_PSII_ of plants under different light conditions. Secondly, we present the development of this model as applied to mung bean (*Vigna radiata* L.) under different light environments, and compare the fitted results with two empirical models which were introduced by Webb *et al*., (1974), Ritchie, (2008), and Smyth *et al*., (2004). Thirdly, we have evaluated these models for the relationship between *Φ*_PSII_ and *I* through comparing the fitted values of *PAR*_sat_ and *ETR*_max_. Finally, we have investigated the impact of the intrinsic characteristics of light-harvesting pigment molecules on *Φ*_PSII_.

## MATERIALS AND METHODS

### Study site and plants

At the end of June 2016, seeds of mung bean *Vigna radiata* [(L.) R. Wilczek] were soaked for 3 h and sowed on potted trays which were filled with matrixes (Scotts Miracle·Gro) consisted of turfy earth, coconut tree branny, pearlite and vermiculite and contained about 2.4 g · kg^-1^ of total nitrogen, 0.95 g · kg^-1^ of P_2_O_5_, 1.27 g · kg^-1^ of K_2_O and about 0.71 g · kg^-1^ of trace elements. Seeds were germinated at the air temperature of 28 °C, the relative humidity of 80 % and light intensity of 125 μmol photons m^-2^ · s^-1^ (12h-day) and grown to the height of 7~10 cm. A total of 44 seedlings (healthy and uniform) were transplanted into 100 mm * 85 mm * 95 mm plastic pots, and divided randomly into four groups with 11 plants per group. They were cultured in phytotron with the day temperature of 25~28°C and the night temperature of 20~24°C. Three light regimes were established: LL (low light with 100 μmol photons m^-2^ · s^-1^), ML (middle light with 220 μmol photons m^-2^ · s^-1^), and HL (high light with 430 μmol photons m^-2^ · s^-1^). Plants were watered every morning up to the full substrate capacity. After 20 days, 5 healthy plants were randomly selected from each light treatment to measure their chlorophyll fluorescence parameters. When the plant height was about 20 cm, the third youngest fully expanded leaf from the top to bottom was used for fluorescence measurements (*n* = 5, *n* – number of replications).

### Chlorophyll fluorescence measurements

Chlorophyll *a* fluorescence was measured on intact leaves using a chlorophyll fluorescence measuring system (Dual PAM-100, Walz, Effeltrich, Germany) with DUAL-E and DUAL-DB measuring heads. The initial level (*F*_0_) of fluorescence was detected after 25 min of dark adaptation. The maximal fluorescence level (*F_m_*) of the dark- and light- 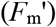 adapted leaves were determined by applying saturating flashes (15.000 μmol photons m^-2^ s^-1^) lasting 1 s, to promote the closure of the PS II reaction centers, according to the method described by Maxwell & Johnson (2000). The fluorescence parameters were calculated as described by van Kooten & Snel (1990) and Klughammer & Schreiber (2008). Maximum quantum efficiency of PS II, *F*_v_/*F*_m_ = (*F*_m_ – *F*_0_)/*F*_m_; non-photochemical quenching, 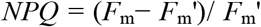. *ETR* was obtained as *ETR*=*α′* × *β′* × *Φ*_PSII_ × *I*, where *α′* is distribution coefficient of absorption light energy by PS II and PSI to be assumed typically 0.5 (Maxwell & Johnson, 2000; Major & Dunton, 2002; Evans, 2009), leaf absorptance (*β*′) is measured using an integrating sphere with a value typically returned of 0.86 (Ehleringer, 1981). Leaves were incrementally exposed to 14 irradiance levels (range 0 to 1450 μmol photons m^-2^ s^-1^) at 30 s intervals.

### Model 1

The photosynthetic electron transport rate via PS II can be described with Eq. 1 (Ye *et al*. 2013a, 2013b),

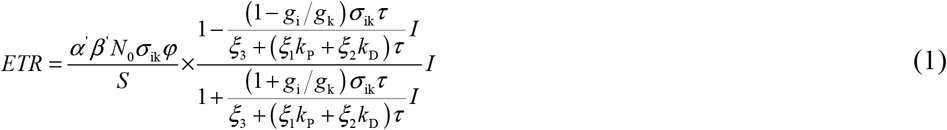

where *φ* is exciton-use efficiency in PS II, *N*_0_ is total photosynthetic pigment molecules of the measured leaf, *S* is the leaf area (m^2^), *g*_i_ and *g*_k_ are the degeneration of energy levels of photosynthetic pigments in the ground state (*i*) and excited state (*k*), respectively. *k*_P_ and *k*_D_ are rates of the photochemical reaction and thermal deactivation, respectively. *ξ*_1_, *ξ*_2_ and *ξ*_3_ are the occupation probability of photochemistry, non-radiation heat dissipation, and fluorescence emission, respectively. *σ*_ik_ is the eigen-absorption cross-section of photosynthetic pigments from the ground state *i* to excited state *k* via light exposure, τ is the average lifetime of the photosynthetic pigments in the lowest excited state *k*.

According to Ye et al., (2013a, 2013b), *σ*_ik_, *τ*, *ξ*_1_, *ξ*_2_, *ξ*_3_, *g*_i_, *g*_k_, *k*_p_ and *k*_D_ (Eq. 1) are inherently specific but have different values depending on species and environmental conditions (e.g. light, temperature, CO_2_ concentration and relative humidity). Therefore, for a given species and at given environmental conditions, we can assume that 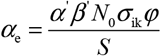 (μmol electrons (μmol photons)^-1^) is the initial slope of light-response curve of electron transport rate, 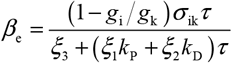 (m^2^ s(μmol photons)^-1^) is the dynamical down-regulation term of PS II, and 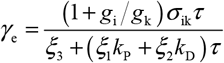 (m^2^ s(μmol photons)^-1^) is the saturation term of photosynthesis. Eq. 1 can be simplified as,

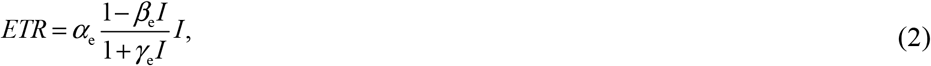

*PAR*_sat_ is calculated from Eq. 3,

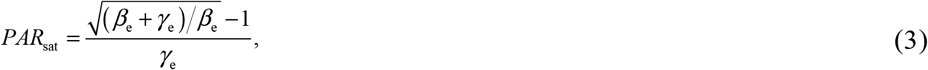

Thus *PAR*_sat_ depends on *σ*_ik_, *τ*, *k*_D_, *k*_P_, *g*_i_, *g*_k_, *γ*_1_, *γ*_2_ and *γ*_3_, but it is independent of *N*_0_.

The maximum value for 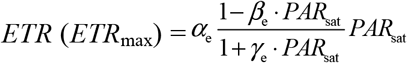, and it can be simplified as,

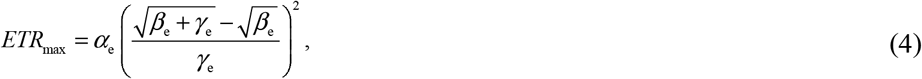

Moreover, compared Eq. 1 with *ETR* =*α′ × β′ × Φ*_PSII_ × *I* (Krall & Edward, 1992), the relationship between *Φ*_PSII_ and *I* is described by Eq. 5,

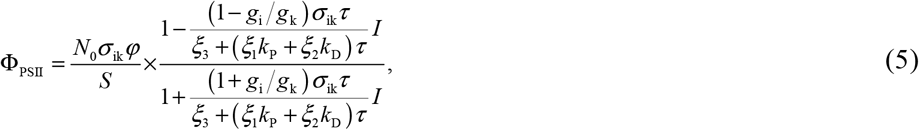

Eq. 5 demonstrates that *Φ*_PSII_ is closely related with intrinsic characteristics of light-harvesting pigment molecules, it does not only depend on *I*, but also on *N*_0_, *σ*_ik_, *τ*, *φ*, *k*_P_, *k*_D_, *g*_i_, *g*_k_, *ξ*_1_, *ξ*_2_ and *ξ*_3_. In particular, Eq. 5 reveals that *Φ*_PSII_ increases with increasing *k*_P_ or *k*_D_, and decreases with increasing *τ*. In addition, Eq. 5 can be simplified as:

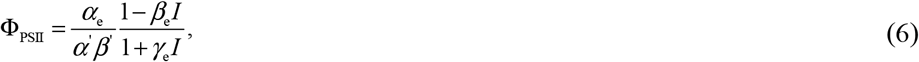

Eq. 6 shows clearly that *Φ*_PSII_ decreases non-linearly with increasing *I* at given environmental conditions (e.g. air temperature, CO_2_ concentration and relative humidity).

In addition, the effective absorption cross-section of light-harvesting pigment molecules 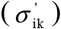 can also be expressed as a function of *I*(Ye *et al*., 2013a, (2013b). Namely,

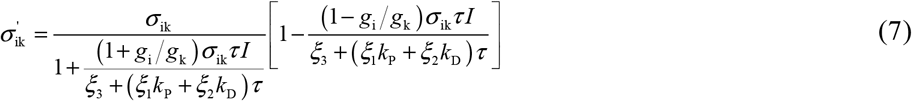

Eq. 7 shows that 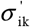 increases with *k*_P_, *k*_D_, *ξ*_1_, *ξ*_2_, *ξ*_3_ and 1/*τ* but decreases with *I*. 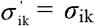 when *I* = 0. As such, the light absorption cross-section is not a constant under any given irradiance (excluding *I* = 0).

Compared Eq. 5 with Eq. 7, the relationship between *Φ*_PSII_ and 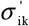 is described by Eq. 8

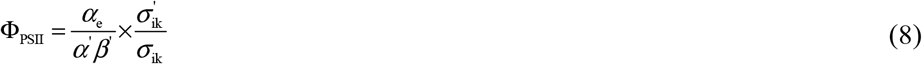

Under given environmental conditions, the values of *α*_e_, *σ*_ik_, *α′* and *β′* are the constants. Therefore, Eq. 8 demonstrates that *Φ*_PSII_ is directly proportional to 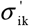 and it changes as a function of 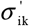.

### Model 2

Effective quantum efficiency of PSII (*Φ*_PSII_) ranges from 0 to 1 (the maximum is not usually higher than 0.85). It has been found experimentally that *Φ*_PSII_ usually follows a simple exponential decay function (Webb *et al*., 1974; Ritchie, 2008 ; Ritchie *et al*., 2010), namely,

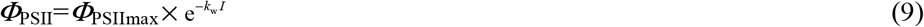

*Φ*_PSIImax_ is the maximum effective quantum efficiency which means the effective quantum efficiency at theoretical zero irradiance, *k*_w_ is a scaling constant, and *I* is the irradiance. The *Φ*_PSIImax_ can be obtained by Eq. 9.

Substituting Eq. 9 into *ETR =α′ × β′ × Φ*_PSII_ × *I* (Krall & Edward, 1992), we get the following expression for *ETR*:

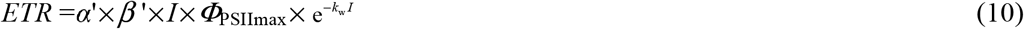

Using Eq. 10, we can calculate saturation irradiance (*PAR*_sat_=1/*k*_w_) and maximum electron transport rate (*ETR*_max_ = *α′ × β′ × PAR*_sat_ × *Φ*_PSIImax_ e^-1^).

Here we take both Eqs. 9 and 10 as model 2.

### Model 3

After Webb *et al*. (1974) introduced an exponential function, Smyth *et al*. (2004) and Silsbe & Kromkamp (2012) used this function to fit light-response curves of *Φ*_PSII_ (Eq. 11).

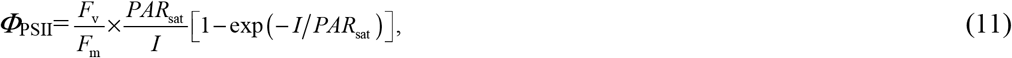

*F*_v_/*F*_m_ is the ‘dark-adapted’ maximum operating efficiency of PS II and *PAR*_sat_ is the saturation irradiance (Smyth *et al*., 2004). The *F*_v_/*F*_m_ and *PAR*_sat_ can be obtained by Eq. 11.

Similarly, substituting Eq. 11 into *ETR =α′ × β′ × Φ*_PSII_ × *I* (Krall & Edward, 1992), we get the following expression for *ETR*,

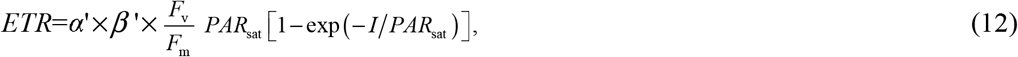

The maximum *ETR* can be calculated by the following formula,

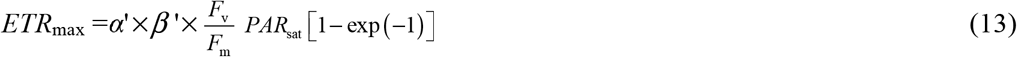

Similarly, here we take Eqs. 12 and 13 as model 3.

### Chlorophyll determination

Leaf discs from control areas were rapidly frozen in liquid nitrogen and ground to powder. Then the chlorophyll (Chl) was extracted with 80% (v/v) acetone and quantified with spectrophotometer (*UVIC0N-930, Kontron Instruments*, Zürich, Switzerland). Chl *a* was determined at wavelength 663 nm, Chl *b* at 646 nm and carotenoids at 470 nm (Lichtenthaler 1987). Total chlorophyll content was determined as described by Porra, Thompson & Kriedermann (1989).

### Statistical analysis

All variables are expressed as mean values (±SE) from five samples for each species. Data were analyzed with one-way analysis of variance (ANOVA) and then the values of *ETR*_max_ and *PAR*_sat_ estimated by three models were compared using a paired-sample *t* test at *α* < 0.05 (*α* - significance level) using the SPSS 18.0 statistical software (SPSS, Chicago, IL). In addition, to compare the advantages and disadvantages of the study models, we took the Akaike’s information criterion (*AIC*) and determination coefficient (*R*^2^) as two indicators to assess the fitting results of the three models. AIC was calculated by reference to Akaike’s method (1973), and *R*^2^ was given directly by SPSS12.5 after fitting the data.

## Results

### Light-response curves of electron transport rate

Representative *ETR-I* curves (fitting the model 1, 2 and 3) for mung bean under three light environments are given in Fig. 1. In LL, *ETR* initially increased (almost linearly) with *I* towards saturation and subsequently, beyond the saturation irradiances exhibited a fast decline, suggesting dynamic down-regulation of PS II or photoinhibition (Fig. 1a). Under ML and HL, beyond *PAR*_sat_, *ETR* values exhibited a little decline with increasing *I* (Fig. 1b and 1c). When the values of *PAR*_sat_ were compared among all the light environments, they ranged from 150 to 417 μmol photons m^-2^ s^-1^. *ETR*_max_ followed a similar trend as *PAR*_sat_ and the values of *ETR*_max_ ranged from 7.36 to 26.04 μmol electrons m^-2^ s^-1^ given the little difference in initial slope of *J-I* curves ranging from 0.456 to 0.586 (Table 2). Moreover, it should be noted that under LL *ETR*_max_ (7.20±0.52 μmol electrons m^-2^ s^-1^), *PAR*_sat_ (149.56±8.81 μmol photons m^-2^ s^-1^) and the initial slope of *ETR-I* curve [ca. 0.456 μmol electrons (μmol photons)^-1^] were lowest among three light environments. This result indicates that the capacity of light energy conversion of plants growing in LL is lower than in ML and in HL.

**Figure 1:**
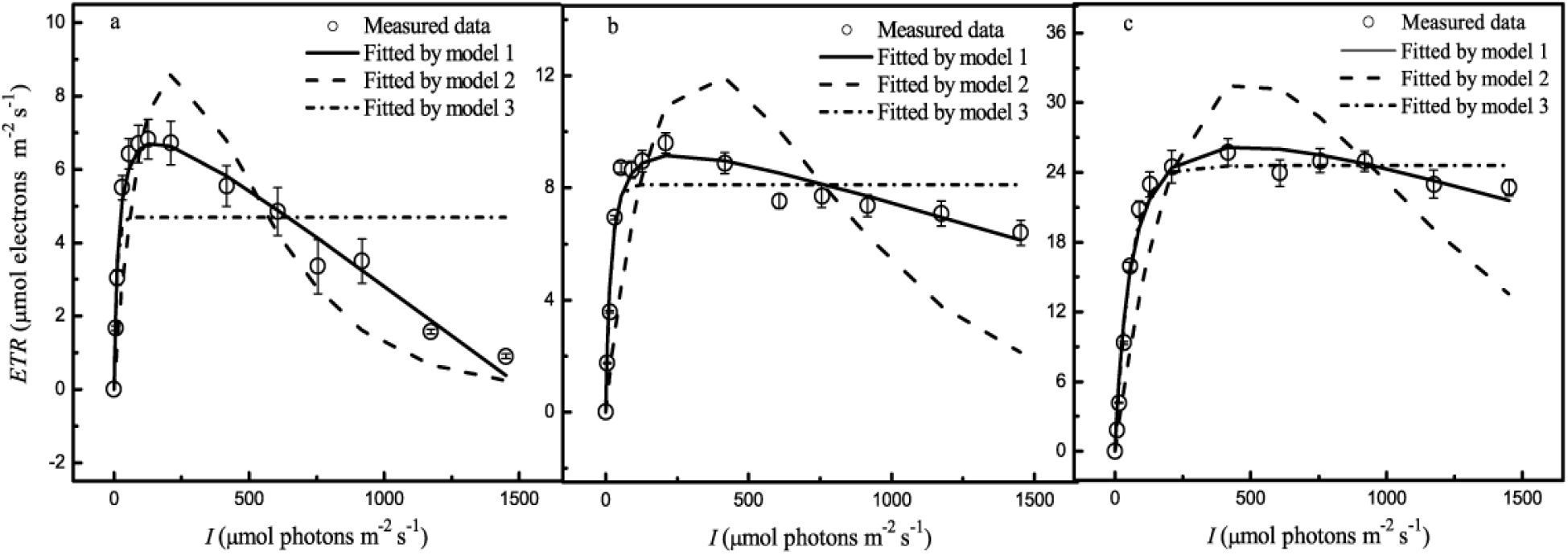
Light-response curves of electron transport rate of mung bean under three light environments (a, LL - 100, b, ML - 200 and c, HL - 340 μmol photons m^-2^ s^-1^) and the curves simulated by models 1, 2 and 3.

**Table 1.**
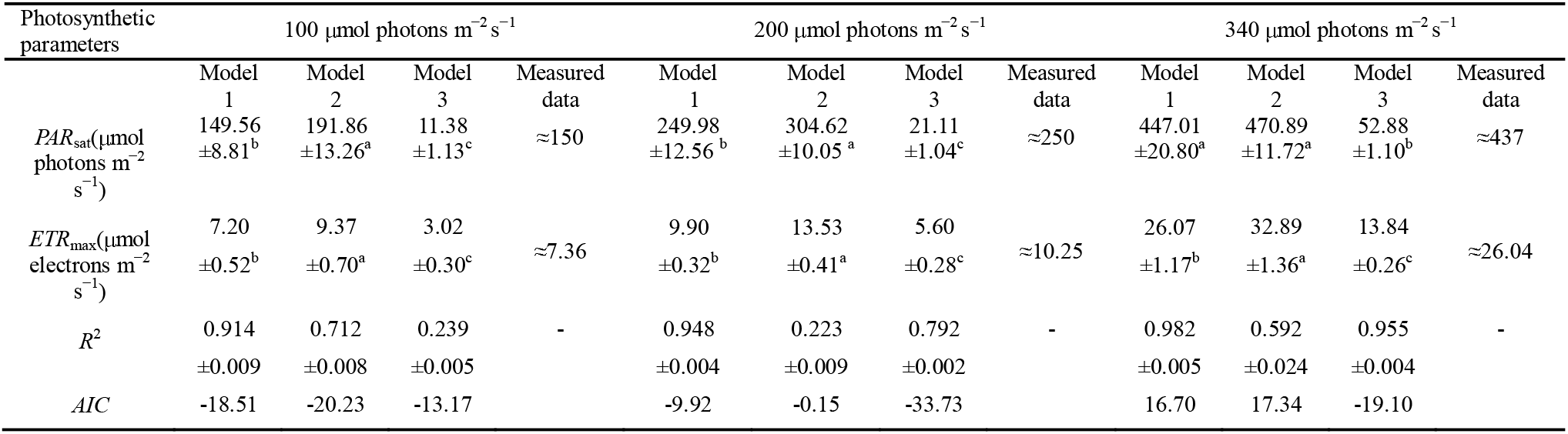
Measured data and results fitted by three models for *ETR-I* curves of mung bean under three light environments. *ETR*_max_ and *PAR*_sat_ estimated by three models were compared using a paired-sample *t* test at α < 0.05 (α - significance level); the values followed by the different superscript letters are significantly different among three models in each light environment. All values indicate the mean ±SE (*n* = 5) except measured data.

**Table 2.** Measured data and results fitted by model 1 for *ETR-I* and *Φ*_PSII_-*I* of mung bean under three light environments. The values of *ETR*_max_ and *PAR*_sat_ estimated by model 1 were compared using a paired-sample *t* test at α < 0.05 (α - significance level); the values followed by the different superscript letters are significantly different in each light environment. All values indicate the mean ± SE (*n* = 5) except measured data.

| Photosynthetic parameters | 100 $\mu\text{mol photons m}^{-2} \text{s}^{-1}$ | | | 200 $\mu\text{mol photons m}^{-2} \text{s}^{-1}$ | | | 340 $\mu\text{mol photons m}^{-2} \text{s}^{-1}$ | | |
| --- | --- | --- | --- | --- | --- | --- | --- | --- | --- |
|  | Model 1 |  |  | Model 1 |  |  | Model 1 |  |  |
| | $\Phi_{PSII-I}$ | $ETR-I$ | Measured data | $\Phi_{PSII-I}$ | $ETR-I$ | Measured data | $\Phi_{PSII-I}$ | $ETR-I$ | Measured data |
| $\Phi_{PSII_{max}}$ | 0.742<br>$\pm 0.016$ | - | - | 0.772<br>$\pm 0.012$ | - | - | 0.756<br>$\pm 0.019$ | - | - |
| $PAR_{sat}$ | 161.19<br>$\pm 12.75^a$ | 149.56<br>$\pm 8.82^a$ | $\approx 150$ | 285.06<br>$\pm 8.62^a$ | 249.98<br>$\pm 12.56^a$ | $\approx 250$ | 396.36<br>$\pm 6.90^a$ | 447.01<br>$\pm 20.80^a$ | $\approx 437$ |
| $ETR_{max}$ | - | 7.20<br>$\pm 0.52$ | $\approx 7.36$ | - | 9.90<br>$\pm 0.32$ | $\approx 10.25$ | - | 26.07<br>$\pm 1.17$ | $\approx 26.04$ |
| $R^2$ | 0.998<br>$\pm 0.005$ | 0.914<br>$\pm 0.009$ | - | 0.996<br>$\pm 0.004$ | 0.948<br>$\pm 0.004$ | - | 0.985<br>$\pm 0.013$ | 0.954<br>$\pm 0.005$ | - |

We used models 1, 2 and 3 to simulate light-response curves of electron transport rate of mung bean under three light environments. Fitted results showed that models 2 and 3 differed markedly from the measured data (Fig. 1). Under three light environments (LL, ML and HL), the coefficients of determination (*R*^2^) for model 1 were 0.914, 0.828 and 0.982; for model 2 *R*^2^ is 0.712, 0.223 and 0.592; for model 3 *R*^2^ is 0.239, 0.792 and 0.955, respectively. Comparing the goodness of fit of the models, model 1, generally, had the best fit and showed greatest *AIC* values.

Under three light environments, the values of *ETR*_max_ and *PAR*_sat_ estimated by model 1 were in agreement with the measured data, whereas model 2 overestimated *ETR*_max_ and *PAR*_sat_, and model 3 greatly underestimated *ETR*_max_ and *PAR*_sat_ (Table 1). *ETR*_max_ and *PAR*_sat_, which are estimated by models 1, 2 and 3, *R*^2^ and *AIC* derived from *ETR-I* curves (Fig. 1) are given in Table 1. Model 1 simulated well *ETR-I* curves, while models 2 and 3 fitted poorly *ETR-I* curves (Fig.1).

### Light-response curves of *Φ*_PSII_

Mung bean under three light environments exhibited a characteristic initial decrease of *Φ*_PSII_ with irradiance (Fig. 1). In LL, the decline of *Φ*_PSII_ differed significantly from HL and decreased more abruptly compared with the other light environments (Fig. 2). It indicates that plants growing in LL had the lowest light energy use efficiency to drive the photochemistry. Moreover, the simulations of the relationship between *Φ*_PSII_ and irradiance by model 1 are in perfect agreement with the experimental data (*R*^2^ ≥ 0.985) (Fig. 2).

**Figure 2:**
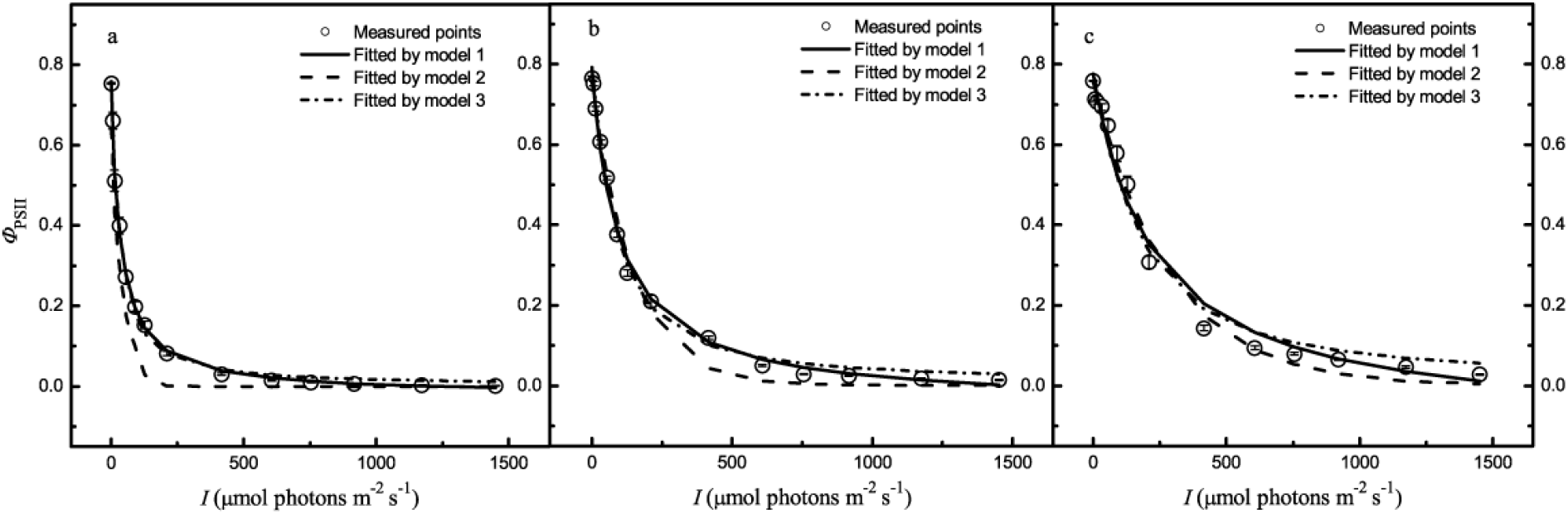
Light-response curves of effective quantum efficiency (*Φ*_PSII_) of mung bean under three light environments (a, LL - 100, b, ML - 200 and c, HL - 340 μmol photons m^-2^ s^-1^) and the curves simulated by models 1, 2 and 3.

In addition, the values of *PAR*_sat_ calculated by Eqs. 3 were not significantly different at a given light intensity (Table 2). *Φ*_PSII_ (and PARsat) calculated by Eqs. 9 (and 10) showed significant differences in each light environment (Table 3). Furthermore, *Φ*_PSII_ (and *PAR*_sat_) calculated by Eqs. 11 and 12 differed significantly in each light environment (Table 4). For example, PARsat estimated by Eqs. 11 and 12 were 11.38 and 24.69 μmol photons m^-2^ s^-1^ in LL, respectively. However, the measured value of PARsat was about 150 μmol photons m^-2^ s^-1^.

**Table 3.**
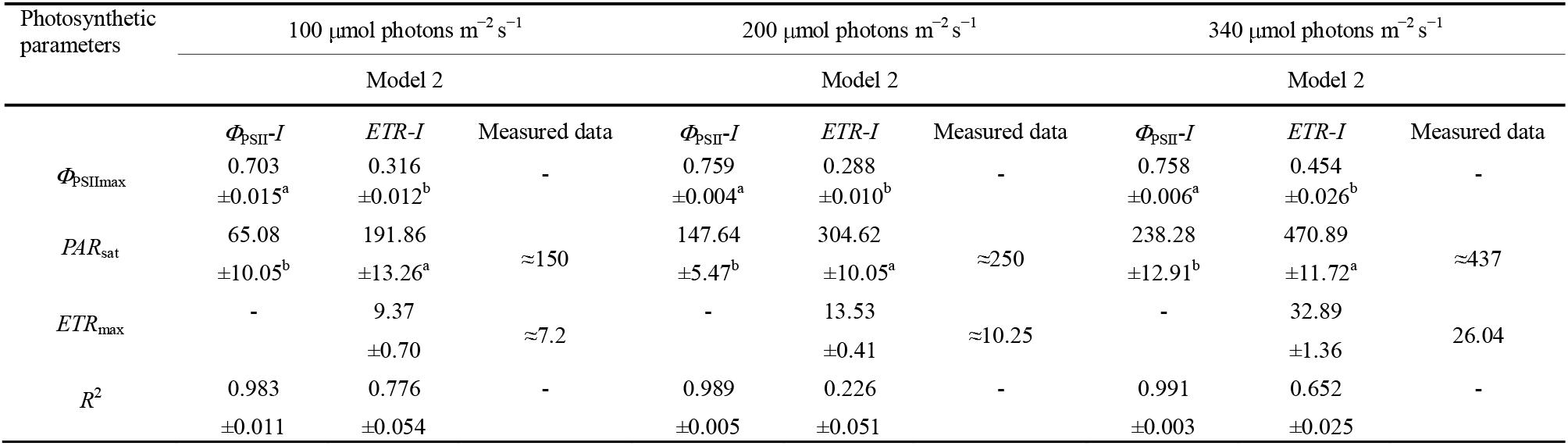
Measured data and results fitted by model 2 for *ETR-I* and *Φ*_PSII_-*I* of mung bean under three light environments. *ETR*_max_, *PAR*_sat_ and *Φ*_PSIImax_ estimated by model 2 were compared using a paired-sample *t* test at α < 0.05 (α - significance level); the values followed by the different superscript letters are significantly different in each light environment. All values indicate the mean ±SE (*n* = 5) except measured data.

**Table 4.** Measured data and results fitted by model 3 for *ETR-I* and *Φ*_PSII_-*I* curves of mung bean under three light environments. *ETR*_max_ and *PAR*_sat_ estimated by model 3 were compared using a paired-sample *t* test at α < 0.05 (α - significance level); the values followed by the superscript letters are significantly different in each light environment. All values indicate the mean±SE (*n* = 5) except measured data.

| Photosynthetic parameters | 100 $\mu\text{mol photons m}^{-2} \text{s}^{-1}$ | | | 200 $\mu\text{mol photons m}^{-2} \text{s}^{-1}$ | | | 340 $\mu\text{mol photons m}^{-2} \text{s}^{-1}$ | | |
| --- | --- | --- | --- | --- | --- | --- | --- | --- | --- |
|  | Model 3 |  |  | Model 3 |  |  | Model 3 |  |  |
| | $ETR-I$ | $\Phi_{PSII}-I$ | Measured data | $ETR-I$ | $\Phi_{PSII}-I$ | Measured data | $ETR-I$ | $\Phi_{PSII}-I$ | Measured data |
| $\Phi_{PSII_{max}}$ | - | - | - | - | - | - | - | - | - |
| $PAR_{sat}$ | 11.38<br>$\pm 1.13^b$ | 24.69<br>$\pm 1.41^a$ | $\approx 150$ | 21.11<br>$\pm 1.04^b$ | 52.61<br>$\pm 1.84^a$ | $\approx 250$ | 52.88<br>$\pm 1.10^b$ | 103.01<br>$\pm 4.49^a$ | $\approx 417$ |
| $ETR_{max}$ | 3.02<br>$\pm 0.030$ | - | $\approx 7.2$ | 5.60<br>$\pm 0.28$ | - | $\approx 10.25$ | 13.84<br>$\pm 0.26$ | - | $\approx 26.04$ |
| $R^2$ | 0.301<br>$\pm 0.048$ | 0.995<br>$\pm 0.008$ | - | 0.795<br>$\pm 0.025$ | 0.996<br>$\pm 0.003$ | - | 0.959<br>$\pm 0.006$ | 0.984<br>$\pm 0.016$ | - |

### Light-response curves of non-photochemical quenching (*NPQ*)

Representative *NPQ-I* curves for mung bean under three light environments are given in Fig. 3. Under LL and ML *NPQ* initially increased (almost linearly) with *I* and subsequently, at the middle irradiances (about 250 μmol photons m^-2^ s^-1^), *NPQ* stabilized and then it increased with *I*. In HL, *NPQ* increased fast when irradiances are beyond 250 μmol photons m^-2^ s^-1^ (Fig. 3c). According to Eqn. (1), plants growing in HL have higher *ETR* than in LL and ML.

**Figure 3:**
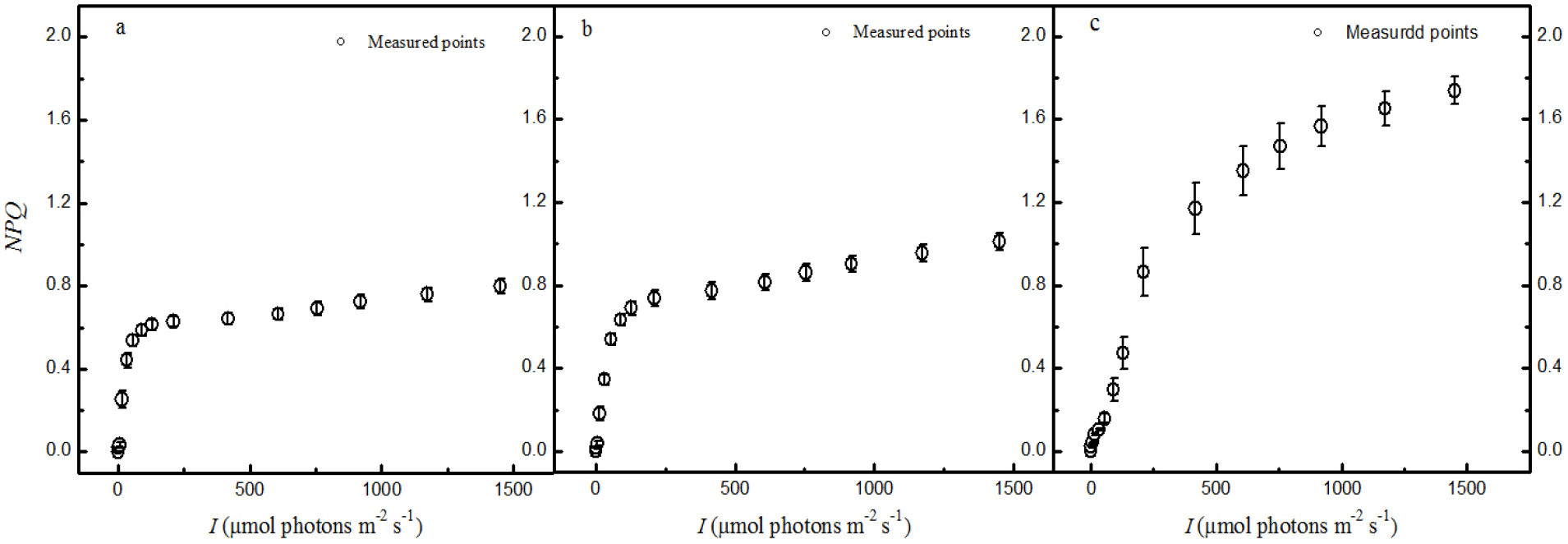
Light-response curves of non-photochemical quenching (*NPQ-I*) of mung bean under three light environments (a, LL - 100, b, ML - 200 and c, HL - 340 μmol photons m^-2^ s^-1^).

### Light-response curves of effective light energy absorption cross-section

The potential for light-harvesting pigments to absorb light energy is reflected by the value of σ_ik_ which increases with the amount of light energy absorbed. For example, in LL, ML and in HL, σ_ik_ for mung bean calculated by 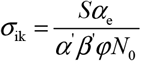 (where *S* is the measured leaf area, *α*_e_ is the initial slope of the light response of photosynthetic electron transport rate, *α′* is the fraction of light absorbed by photosystem II, *β′* is leaf absorptance, *φ* is excitation efficiency of PS II, i.e. for charge separation of P680) is (7.62±0.27) × 10^-21^ m^2^, (9.85±0.46) × 10^-21^ m^2^ and (11.65±0.68) × 10^-21^ m^2^, respectively; these values are all significantly different form one another (*n* = 5, one-way ANOVA followed by Tukey’s test, *α* < 0.05). Eq. 7 shows that 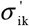 increases with *k*_p_ or *k*_D_, but decreases (non-linearly) with increasing values of *I*, and/or *τ*. Therefore, the competition for each exciton amongst photochemistry, heat dissipation and fluorescence emission directly affects the effective light energy absorption cross-section of light-harvesting pigments 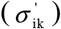 (Fig. 4). At a given irradiance in 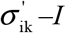 curves, the higher 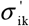 is, the more light energy it absorbs. For example, in Fig. 4 at *I*= 600 μmol photons m^-2^ s^-1^ 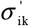 is 0.12 × 10^-21^ m^2^, 0.26 × 10^-21^ m^2^ and 0.36 × 10^-21^ m^2^ under LL, ML and HL. It indicated that under HL mung bean had a higher ability to absorb light energy than under LL and ML.

**Figure 4:**
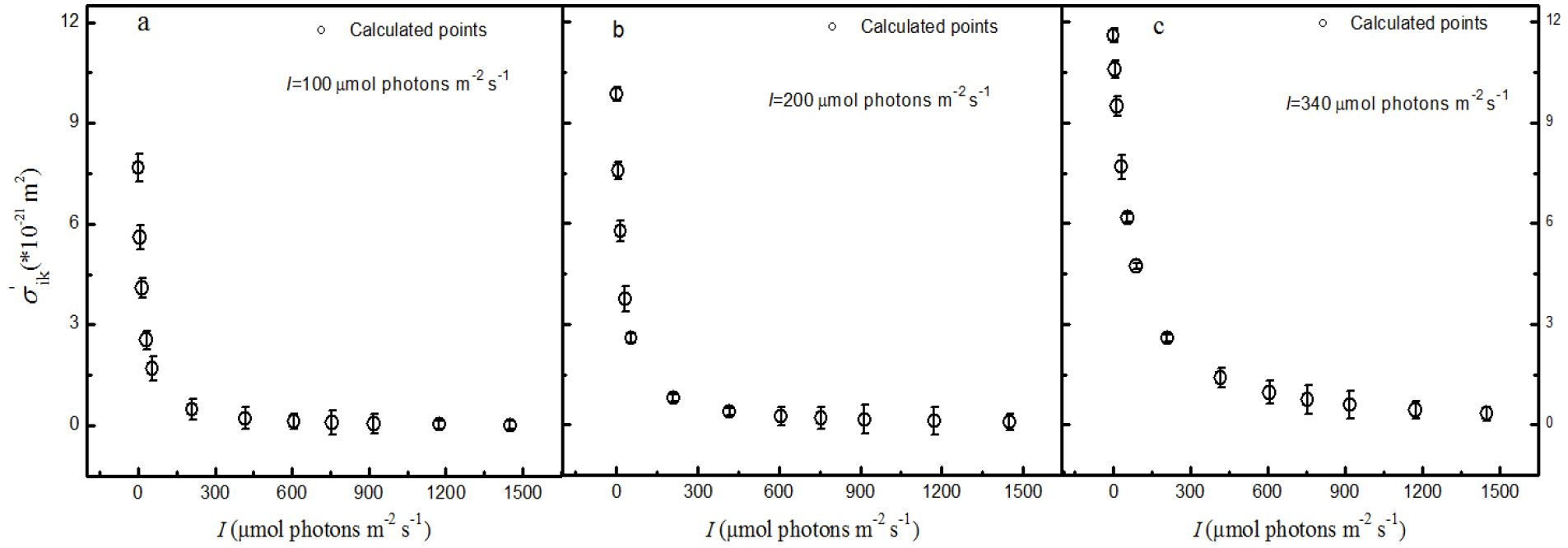
Light-response curves of effective light energy absorption cross-section 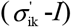 of mung bean under three light environments (a, LL - 100, b, ML - 200 and c, HL - 340 μmol photons m^-2^ s^-1^).

## Discussion

We have built a new model of the relationship between *Φ*_PSII_ and *I* based on light absorption and energy transfer to the reaction centers via photosynthetic pigments. It includes all relevant processes involving light energy absorption and conversion and transfer to the reaction centers of PS II (Ye *et al*., 2013a, 2013b).

The present study was focused on the relationships between *Φ*_PSII_, *ETR*, *σ*_ik_ and *I* in mung bean seedlings growing in one of three light environments. Our main results were the Eq. 1 and 2 used to determine *ETR* in function of *I* and explain the interdependence between *ETR* and biophysical parameters, such as 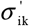. What is most important, we found that the *Φ*_PSII_ decrease is caused by the reduction of 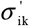 with increasing I. This finding is of fundamental importance to understand the molecular and biophysical mechanisms of variation in *Φ*_PSII_ under the changing light environment. In our study *Φ*_PSII_ not only depends on *N*_0_, *σ*_ik_ and *φ* when environmental factors (e.g. irradiance, CO_2_ concentration and temperature) are constant, but it depends also on *τ*, *k*_P_, *k*_D_, *g*_i_, *g*_k_, *ξ*_1_, *ξ*_2_ and *ξ*_3_, specially on *τ*, *k*_P_ and *k*_D_ which are closely associated with processes of light energy absorption, excitons production and excitation energy transfer to reaction centers. In Eq. 5 *Φ*_PSII_ increases with *k*_P_ (Fig. 1) and *k*_D_ (Fig. 3) when they increase with light intensity, but *Φ*_PSII_ decreases with *τ*.

We used models 1, 2 and 3 to simulate *ETR-I* and *Φ*_PSII_-*I* curves of mung bean in three light environmental conditions. There was evidence that model 2 and model 3 did not satisfactorily fit *ETR-I* and *Φ*_PSII_-*I* curves. Generally, the values of *PAR*_sat_ and *ETR*_max_ estimated by model 2 were higher than the measured values (Table 2), while the values of *PAR*_sat_ and *ETR*_max_ estimated by model 3 were lower than the measured values (Table 2). These results were similar to Laws *et al*. (2002), Smyth *et al*. (2004), Silsbe & Kromkamp (2012). The values of *PAR*_sat_ and ETR_max_ estimated by model 1 were very close to the measured values (Table 2). Thus, we concluded that models 2 and 3 simulate poorly *ETR-I* and *Φ*_PSII_-*I* curves, especially under high level of actinic light inducing dynamic down-regulation of PS II.

Mathematical models are not only useful to reproduce and explain the observed plants behavior, but also to make predictions and attempt to answer more fundamental questions. For example, model 1 (Eq. 5) forecasts increases in *Φ*_PSII_ with *k*_P_ and *k*_D_, and decreases with increasing *τ*. Our results confirm that *Φ*_PSII_ increases with *k*_P_ and *k*_D_. In HL *k*_P_ (reflected in *ETR*) and *k*_D_ (reflected in *NPQ*) (Table 2 and Fig. 3) were higher compared with LL and ML. What is of great importance, based on Eq. 5 or 8, the reason that *Φ*_PSII_ decreases with increasing *I* is the decrease in 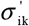. In this study, in HL, mung bean shows the most abrupt decrease in 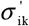 compared with LL and ML (Fig. 4).

So far we have deliberately excluded that the environmental factors influence *Φ*_PSII_ to focus on properties of photosynthetic pigments and PSII which may have a significant effect on *Φ*_PSII_. With the model presented here, it is straightforward to include equations representing *ETR-I* and *Φ*_PSII_-*I* which are associated with light energy absorption, excitons production and its transfer to photochemical reaction centers, which may answer why LL mung bean has the lowest *Φ*_PSII_, *ETR*_max_ and *PAR*_sat_ values, and also explain why its *ETR*s decline fastest compared with ML and HL plants. If the model of the relationship between *Φ*_PSII_ and *I* is combined with Eq. 1 (Ye *et al*. 2013a), much information on physical and biochemical parameters of photosynthetic pigments could be obtained. Additionally, the model of the relationship between *Φ*_PSII_ and *I* indicates *Φ*_PSII_ to be proportional to the product of *N*_0_, *φ* and *σ*_ik_ besides *τ*, *k*_P_, *k*_D_, *g*_i_, *g*_k_, *ξ*_1_, *ξ*_2_ and *ξ*_3_. Therefore, plants with a higher product of *N*_0_, *φ* and *σ*_ik_ have a higher initial value of *Φ*_PSII_.

In conclusion, the proposed model provides the means to predict and simulate the *Φ*_PSII_-*I* curves. The *Φ*_PSII_ decrease with increasing *I* stems from the decrease in effective light energy absorption cross-section of photosynthetic pigments (see Ye *et al*., 2013a, 2013b). When the presented model is combined with the mechanistic model of *ETR-I*, they could become an effective tool towards identifying novel mechanistic properties by which plants modify their light harvesting properties and show photoacclimation (Dubinsky & Stambler, 2009), photoprotection (Takahashi & Badger, 2011; Niyogi & Truong, 2013), dynamic down-regulation of PSII (Ralph & Gademann, 2005) or photoinhibition (Govdjee, 2002) in response to environmental stressors. A cutting-edge next step will be to investigate more species adapted to different environmental conditions along geographical gradients to distinguish between genetic and environmental factors effects on *Φ*_PSII_.

## Acknowledgements

This research was supported by the Natural Science Foundation of China (Grant No. 31560069) and the Natural Science Foundation of Jiangxi Province (Grant No. 20142BAB20402).

## References

Baker NR. 2008. Chlorophyll fluorescence: A probe of photosynthesis in vivo. Annual of Review Plant Biology 59, 89–113.

Buckley TN, Farquhar GD. 2004. A new analytical model for whole-leaf potential electron transport rate. Plant, Cell and Environment 27, 1487–1502.

Córdoba J, Molina-Cano J-L, Martínez-Carrasco R, Morcuende R, Pérez P. 2016. Functional and transcriptional characterization of a barley mutant with impaired photosynthesis. Plant Science 244, 19–30.

Dubinsky Z, Stambler N. 2009. Photoacclimation processes in phytoplankton: mechanisms, consequences, and applications. Aquatic Microbial Ecology 56, 163–176.

Ehleringer JR. 1981. Leaf absorptances of Mohave and Sonoran deserts plants. Oecologia 49, 366–370.

Evans JR. 2009. Potential error in electron transport rates calculated from chlorophyll fluorescence as revealed by multilayer leaf model. Plant and Cell Physiology 50, 698–706.

Fryer MJ, Andrews JR, Oxborough K, Blowers DA, Baker NR. 1998. Relationship between CO_2_ assimilation, photosynthetic electron transport, and active O_2_ metabolism in leaves of maize in the field during periods of low temperature. Plant Physiology 116, 571–580.

Genty B, Briantais JM, Baker NR. 1989. The relationship between the quantum efficiency of photosynthetic electron transport and quenching of chlorophyll fluorescence. Biochimica et Biophysica Acta 990, 87–92.

Govindjee 2002. A role for a light-harvesting antenna complex of photosystem II in photoprotection. Plant Cell 14, 1663–1668.

Kalaji HM, Jajoo A, Oukarroum A, Brestic M, Zivcak M, Samborska IA, Cetner MD, Łukasik I, Goltsev V, Ladle RJ. 2016. Chlorophyll a fluorescence as a tool to monitor physiological status of plants under abiotic stress conditions. Acta Physiologiae Plantarum 38, 102.

Klughammer C, Schreiber U. 2008. Complementary PS II quantum yields calculated from simple fluorescence parameters measured by PAM fluorometry and the saturation pulse method. PAM Application Notes 1, 11–14

Krall JP, Edward GE. 1992. Relationship between photosystem II activity and CO_2_ fixation in leaves. Physiologia Plantarum 86, 180–187.

Krause GH, Weis E. 1984. Chlorophyll fluorescence as a tool in plant physiology. II. Interpretation of fluorescence signals. Photosynth Research 5, 139–157

Laws E, Sakshaug E, Babin M, Dandonneau Y, Falkowski P, Geider R, Legendre L, Morel A, Sondergaard M, Takahashi M, Williams PJ. 2002. Photosynthesis and primary productivity in marine ecosystems: Practical aspects and application of techniques. JGOFS Report No. 36.

Lichtenthaler HK. 1987. Chlorophyll and carotenoids – pigments of photosynthetic biomembranes. Method in Enzymology 148, 350–382.

Majláth I, Darko E, Palla B, Nagy Z, Janda T, Szalai G. 2016. Reduced light and moderate water deficiency sustain nitrogen assimilation and sucrose degradation at low temperature in durum wheat. Journal of Plant Physiology 191, 149–158.

Major KM, Dunton KH. 2002. Variations in light-harvesting characteristics of the seagrass, *Thalassia testudinum*: evidence for photoacclimation. Journal of Experimental Marine Biology and Ecology 275, 173–189.

Maxwell K, Johnson NG 2000. Chlorophyll fluorescence—a practical guide. Journal of Experimental Botany 51, 659–668.

Moin M, Bakshi A, Saha A, Kumar MU, Reddy AR, Rao KV, Siddiq EA, Kirt PB. 2016. Activation tagging in Indica rice identifies ribosomal proteins as potential targets for manipulation of water-use efficiency and abiotic stress tolerance in plants. Plant, Cell and Environment 39, 2440–2459.

Niyogi KK and Truong TB. 2013. Evolution of flexible non-photochemical quenching mechanisms that regulate light harvesting in oxygenic photosynthesis. Current Opinion in Plant Biology 16, 307–314.

Oxborough K. 2004. Imaging of chlorophyll a fluorescence: theoretical and practical aspects of an emerging technique for the monitoring of photosynthetic performance. Journal of Experimental Botany 55, 1195–1205.

Panitchayangkoon G, Hayes D, Fransted KA, Caram JR, Harel E, Wen J, Blankenship RE, Engel GS. 2010. Long-lived quantum coherence in photosynthetic complexes at physiological temperature. Proceedings of National Academy of Sciences of the United States of America 107, 12766–12770.

Pavlovic A, Slovakova L, Pandolfi C, Mancuso S. 2011. On the mechanism underlying photosynthetic limitation upon trigger hair irritation in the carnivorous plant venus flytrap (*Dionaea muscipula* Ellis). Journal of Experimental Botany 62, 1991–2000.

Porra RJ, Thompson WA, Kriedermann PE. 1989. Determination of accurate extinction coefficient and simultaneous equations for essaying chlorophylls a and b extracted with four different solutions: verification of the concentration of chlorophyll standards by atomic absorption spectroscopy. Biochemica and Biophysica Acta 975, 384–394.

Ralph PJ, Gademann R. 2005. Rapid light curves: A powerful tool assess photosynthetic activity. Aquatic Botany 82, 222–237.

Rascher U, Liebieg M, Lüttge U. 2000. Evaluation of instant light-response curves of chlorophyll fluorescence parameters obtained with a portable chlorophyll fluorometer on site in the field. Plant, Cell and Environment 23, 1397–1405.

Renk H, Ochocki S, Kurzyk S. 2000. *In situ* and *simulated in situ* primary production in the Gulf of Gdańsk. Oceanologia 42, 263–282.

Richter M, Renger T, Knorr A. 2008. A Block equation approach to intensity dependent optical spectra of light harvesting complex II. Photosynthesis Research 95, 119–127.

Ritchie RJ. 2008. Fitting light saturation curves measured using modulated fluorometry. Photosynthesis Research 96, 201–215.

Ritchie RJ, Bunthawin S. 2010. The use of pulse amplitude modulation (PAM) fluorometry to measure photosynthesis in a cam orchid, *Dendrobium* spp. (D. cv. *Viravuth pink*). International Journal of Plant Science 171, 575–585.

Robakowski P. 2005. Susceptibility to low-temperature photoinhibition in three conifers differing in successional status. Tree Physiology 25, 1151–1160.

Sarovar M, Ishizaki A, Fleming GR, Whaley KB. 2010. Quantum entanglement in photosynthetic light-harvesting complexes. Nature Physics 6, 462–467.

Silsbe GM, Kromkamp JC. 2012. Modeling the irradiance dependency of the quantum efficiency of photosynthesis. Limnology and Oceanography: Methods 10, 645–652.

Smyth TJ, Pemberton KL, Aiken J, Geider RJ. 2004. A methodology to determine primary production and phytoplankton photosynthetic parameters from Fast Repetition Rate Fluorometry. Journal of Plankton Research 26, 1337–1350.

Suggett DJ, Prasil O, Borowitzka MA. 2010. Chlorophyll a fluorescence in aquatic sciences: methods and applications (developments in applied phycology). Springer, London, pp 323.

Takahashi S, Badger MR. 2011. Photoprotection in plants: a new light on photosystem II damage. Trends in Plant Science 16, 53–59.

van der Tol C, Berry JA, Campbell PKE, Rascher U. 2014. Models of fluorescence and photosynthesis for interpreting measurements of solar-induced chlorophyll fluorescence, Journal of Geophysical Research: Biogeosciences 119, 2312–2327.

van Kooten O, Snel JFH. 1990. The use of chlorophyll fluorescence nomenclature in plant stress physiology. Photosynthesis Research 25, 147–150.

Ye ZP, Robakowski P, Suggett DJ. 2013a. A mechanistic model for the light response of photosynthetic electron transport rate based on light harvesting properties of photosynthetic pigment molecules. Planta 237, 837–847.

Ye ZP, Suggett DJ, Robakowski P, Kang HJ. 2013b. A mechanistic model for the photosynthesis-light response based on the photosynthetic electron transport of PSII in C_3_ and C_4_ species. New Phytologist 199, 110–120.

